# *In vivo* Locus Coeruleus activity while awake is associated with REM sleep quality in healthy older individuals

**DOI:** 10.1101/2023.02.10.527974

**Authors:** Ekaterina Koshmanova, Alexandre Berger, Elise Beckers, Islay Campbell, Nasrin Mortazavi, Roya Sharifpour, Ilenia Paparella, Fermin Balda, Christian Berthomier, Christian Degueldre, Eric Salmon, Laurent Lamalle, Christine Bastin, Maxime Van Egroo, Christophe Phillips, Pierre Maquet, Fabienne Collette, Vincenzo Muto, Daphne Chylinski, Heidi IL Jacobs, Puneet Talwar, Siya Sherif, Gilles Vandewalle

## Abstract

The locus coeruleus (LC) is the primary source of norepinephrine (NE) in the brain, and the LC-NE system is involved in regulating arousal and sleep. It plays key roles in the transition between sleep and wakefulness, and between slow wave sleep (SWS) and rapid eye movement sleep (REMS). However, it is not clear whether the LC activity during the day predicts sleep quality and sleep properties during the night, and how this varies as a function of age. Here, we used 7 Tesla functional Magnetic Resonance Imaging (7T fMRI), sleep electroencephalography (EEG) and a sleep questionnaire to test whether the LC activity during wakefulness was associated with sleep quality in 52 healthy younger (N=33; ~22y; 28 women) and older (N=19; ~61y; 14 women) individuals. We find that, in older, but not in younger participants, higher LC activity, as probed during an auditory mismatch negativity task, is associated with worse subjective sleep quality and with lower power over the EEG theta band during REMS (4-8Hz), which are two sleep parameters significantly correlated in our sample of older individuals. The results remain robust even when accounting for the age-related changes in the integrity of the LC. These findings suggest that the activity of the LC may contribute to the perception of the sleep quality and to an essential oscillatory mode of REMS, and that the LC may be an important target in the treatment of sleep disorders and age-related diseases.

## Introduction

Sleep is essential to health. Insufficient or poor sleep impacts cognitive, attentional and learning abilities at all ages (1–3), while in the long run, it also increases the risk of developing diabetes (4), cardiovascular diseases (5,6), mood disorders (7,8) and neurodegeneration (9). Sleep quality declines over the adult lifespan, with an elevated rate of sleep complaints and sleep disorders in aging individuals (10). These could arguably contribute to the higher prevalence of psychiatric and neurological diseases reported at older ages (11). Here, we posit that the link between sleep quality and aging arises, at least partly, from the locus coeruleus (LC), a small nucleus in the brainstem.

The LC constitutes the primary source of norepinephrine (NE) in the central nervous system and sends ubiquitous monosynaptic projections to almost all brain areas. The LC-NE system plays a primary role in many aspects of brain functions, including the maintenance of wakefulness (12), sleep onset, and the alternation between slow wave sleep (SWS) and rapid eye movement sleep (REMS) (13,14). The integrity of the LC is progressively altered over adulthood. The LC contrast measured using magnetic resonance imaging (MRI) and considered to reflect the neuronal density of the LC, increases up to about 60 years of age and then declines afterwards (Liu et al., 2019; Shibata et al., 2006). The LC is also one of the first brain sites to show, in otherwise healthy individuals, pretangle tau material that is later co localized with insoluble tau tangles, and synuclein inclusions, which are the hallmarks of the neuropathology of Alzheimer’s and Parkinson’s diseases (AD, PD), respectively (15,16). Importantly, degeneration of the LC neurons contributes to the pathophysiology of REMS behavioral disorder, a preclinical PD condition (17). It is therefore plausible that the age-related changes in LC integrity affect its functions and contribute, in turn, to the age-related alterations in sleep quality.

Despite the strong link between the LC and sleep, most of the research on this topic was conducted on animal models. There is conflicting evidence in early studies on the consequences of the LC lesions on sleep-wake states, and more recent research have demonstrated that the inhibition of the LC reduces time spent in wakefulness and its activation leads to sleep-to-wake transitions (18). Furthermore, the duration of REMS and the probability REMS – non-REMS (NREMS) transitions did not directly depend on the LC inhibition / stimulation, which implies a modulatory involvement of the LC to REMS rather than a direct contribution to its genesis. Importantly, in human research, poor structural integrity of the LC (assessed with dedicated-MRI-derived LC contrast) was recently linked to a higher number of nocturnal awakenings in older cognitively unimpaired individuals, especially in the presence of AD biomarkers (19). Translation of animal findings to human beings may not be straightforward (12), and to date, there is no report of an *in vivo* assessment of the LC functioning in relation to sleep characteristics. This is likely due to the deep position and the small size of the LC - ~15 mm long, ~2.5 mm diameter, ~50.000 neurons (20). Part of these limitations are being lifted by the advent of ultra-high field 7 Tesla (7T) MRI which provides a higher signal-to-noise ratio and a higher resolution than most commonly used 3T MRI.

The difficulty of imaging the LC also arises from the fact that the tonic activity of the LC, which is state dependent, is reduced during SWS compared to wakefulness and (almost) absent during REMS, while its highest during wakefulness fluctuating with the level of attentiveness (21,22). Asides from their tonic mode of activity, the LC neurons can also function following a phasic mode (23). Phasic bursts happen in response to salient stimuli, and performance to an attentional task follows inverted-U shaped curve depending on the interplay between phasic and tonic discharge activity (23). One could therefore argue that variability in LC function as captured in an attentional task reflects the variability in the processes modulated by the LC, including sleep.

Here, we tested whether the LC activity probed during wakefulness, is associated with the quality of sleep in healthy younger and older late middle-aged individuals, respectively aged 18 to 30 and 50 to 70 y. Participants’ brain activity was recorded in a 7T MRI scanner while they completed a mismatch negativity attentional task, which mimics novelty and salience detection and is known to elicit LC activity (24). Subjective sleep quality was assessed by a questionnaire, together with objective sleep measures as extracted from electroencephalogram (EEG) recordings during a nocturnal sleep session. We hypothesized that higher activity of the LC during wakefulness would be associated with worse subjective and objective sleep quality.

## Results

Fifty-two healthy participants with no history of cognitive and sleep disorders completed the study, including 33 young adults (22.3 ± 3.2 y; 28 women) and 19 late middle-aged individuals (61.05 ± 5.3 y; 14 women) (**Table 1**). They first completed a structural 7T MRI session, which served as habituation to the MR environment, and allowed reconstruct a high-resolution whole-brain image as well as a dedicated LC specific image (**Figure 1A**). The latter was used to create individual LC-masks in each participant’s brain space that were averaged into a group-wise LC-mask in a standardized brain space (see methods). Participants were requested to sleep regularly prior to completing a fMRI session (see methods) in the morning, 2 to 3h after wake-up time, during which they performed an auditory oddball task (**Figure 1C**) (24). Participants further provided a subjective evaluation of their habitual sleep quality using a validated questionnaire: the Pittsburgh Sleep Quality Index (PSQI, (25)). Their habitual baseline sleep was recorded in-lab under EEG to extract our main objective sleep features of interest spanning some of the most canonical characteristics of sleep (**Figure 1B**): sleep onset latency, related to sleep initiation; sleep efficiency (ratio between sleep time and time in bed), to assess overall sleep quality and continuity; REMS percentage, to reflect the global architecture of sleep; slow wave energy (SWE) during SWS (cumulated overnight 0.5-4Hz EEG power), an accepted marker of sleep need related to the intensity of SWS (26); and the cumulated overnight power over the theta band of the EEG (4-8 Hz) during REMS, associated with REMS intensity over its most typical oscillatory activity.

**Figure 1.**
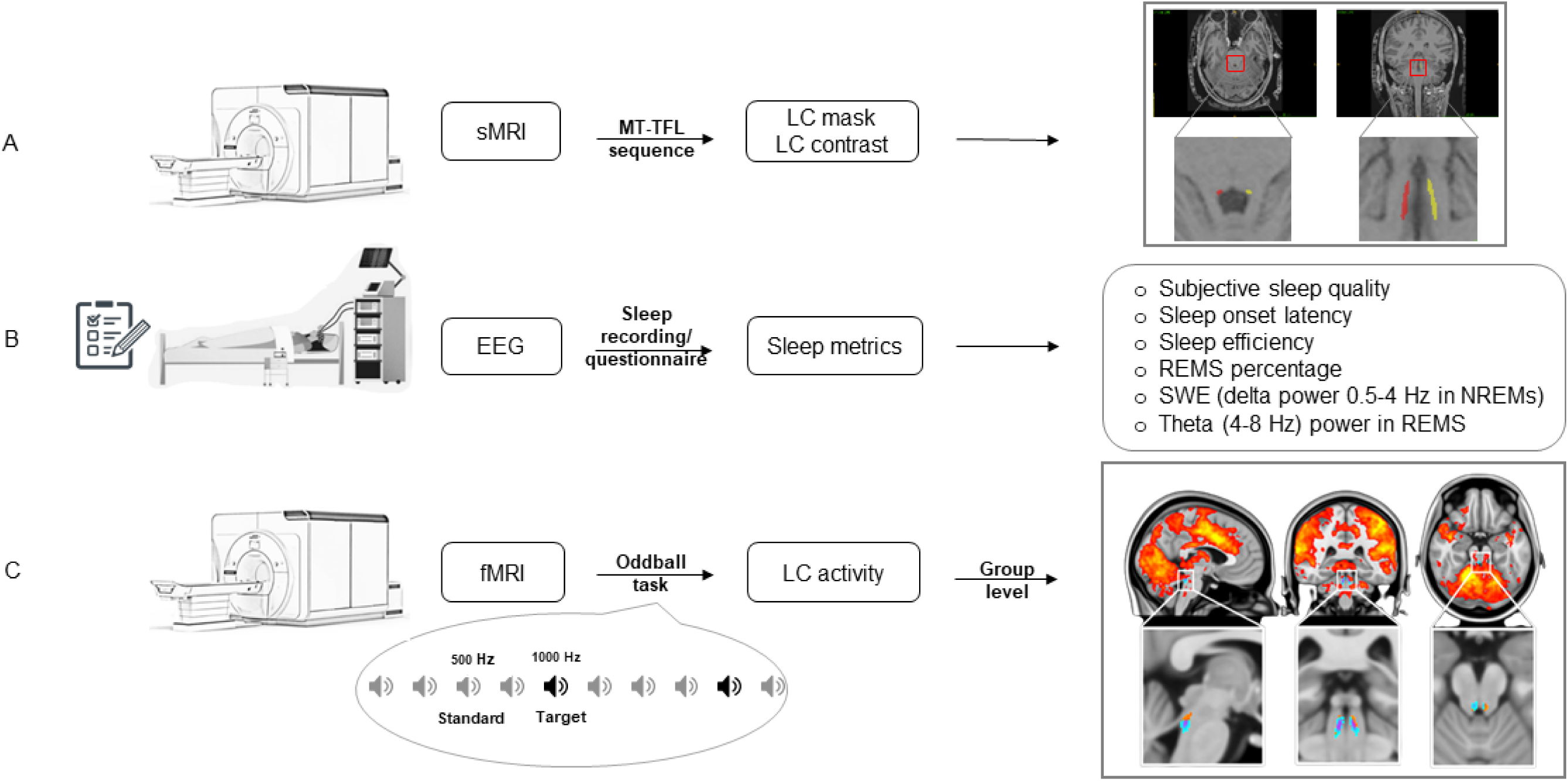
Overview of the study protocol. (A) The volunteers completed a structural 7T MRI (sMRI) session including a sequence for the segmentation of the LC. The latter was used to create individual LC-mask in each participant’s brain space as shown in a representative subject (red: left LC; yellow: right LC) and to compute the LC contrast, reflecting the structural integrity of the LC. (B) Participants’ habitual baseline sleep data was recorded overnight in-lab under EEG before the fMRI session to extract our main objective sleep features of interest. They further provided a subjective evaluation of their habitual sleep quality using a validated questionnaire. (C) After the baseline night, participants underwent a fMRI session during which they completed an auditory oddball task. Brain responses to the deviant tones are displayed as in (27) over the group average brain structural image [top row; p<.001 uncorrected, right color legend refers to t-values between 3.26 – red - and 8 – yellow] (top) and only over the group-wise of the LC built based on individual LC masks (bottom ro; p<.05 FWE corrected).

**Table 1:** Characteristics of the study sample.

|  | **Late middle-aged**<br>**(n = 19)** |  |  |  | **Young**<br>**(n = 33)** |  |  |  |  |
| --- | --- | --- | --- | --- | --- | --- | --- | --- | --- |
|  | **Mean** | **SD** | **Min** | **Max** | **Mean** | **SD** | **Min** | **Max** | **p-value** |
| *Age* | 61 | 5.3 | 53 | 70 | 22 | 2.8 | 18 | 29 | 0.000 |
| *BMI (kg/m<sup>2</sup>)* | 24.9 | 3.52 | 19.4 | 30.9 | 21.96 | 3.11 | 17.2 | 28.4 | 0.005 |
| *Education (years)* | 14.58 | 2.63 | 9 | 19 | 14.5 | 2.28 | 12 | 20 | 0.81 |
| *BDI* | 5.58 | 4.05 | 0 | 14 | 6.73 | 4.2 | 0 | 20 | 0.26 |
| *BAI* | 3.16 | 3.13 | 0 | 9 | 3.94 | 2.9 | 0 | 11 | 0.33 |
| *ESS* | 5.52 | 3.82 | 0 | 13 | 7 | 3.86 | 0 | 14 | 0.15 |
| *PSQI* | 3.95 | 2.22 | 0 | 8 | 4.61 | 1.97 | 1 | 9 | 0.24 |
| *Sex (F - M)* | 14 F – 5 M |  |  |  | 28 F – 5 M |  |  |  | 0.15 |

The education level is expressed in the number of years of formal schooling. BMI stands for Body Mass Index. BDI score, BAI score, ESS score and PSQI respectively stand for Beck Depression Inventory score (55), Beck Anxiety Inventory score (54), Epworth Sleepiness Scale score (71) and Pittsburgh Sleep Quality Index (25). F: Female and M: Male.

As already reported in this sample (27), we found significant activations within the bilateral (though more left lateralized) rostral part of the LC for the detection of the target sound (**Figure 1C**; p <.05, FWE corrected for multiple comparisons over the group-wise LC-mask), supporting a robust LC activation during the oddball task. Individual estimates of LC activity were then extracted within the entire individual LC masks in the participant brain space, for higher accuracy, and contrasted against the sleep metrics of interest.

Our first statistical test asked whether habitual subjective sleep quality was related to the activity of the LC. A generalized linear mixed model (GLMM) with subjective sleep quality as the dependent variable found a main effect of LC activity (**p = 0.017**) and age group (**p = 0.046**), as well as a significant LC-activity-by-age-group interaction (**p = 0.006**), whilst controlling for sex and BMI (**Table 2**). Post-hoc tests revealed that higher LC activity was associated with worse subjective sleep quality in the older (t = 2.81, **p = 0.007**), but not in the younger group (t = −0.77, P = 0.45) (**Figure 2A**). We computed the same GLMM using the mean activity of the left and right LC separately to assess whether there is a lateralized association. We obtained similar statistical outputs, though more prominently when focusing on the left LC (main effect of LC activity – left: F = 5.1, **p= 0.03,** R^2*^ = 0.1 – right: F = 2.46, p= 0.12; left and right LC-activity-by-age-group interactions: F> 5, **p≤ 0.03**). Both with the left and the right LC activity, post-hoc tests yielded a significant correlation in the older (t > 2, **p≤ 0.04**), but not in the younger group (−0.9 < t < 0, p > 0.39). Furthermore, since we previously reported in our sample a significant age group difference in LC integrity – as indexed by its contrast (see methods and (27)) - we added LC contrast as a covariate in the GLMM which yielded the same main effects of LC activity and LC-activity-by-age-group interaction (**Table S1**). Interestingly, the GLMM yielded a significant main effect of LC contrast (F = 5.34; **p=0.025,** R^2*^ = 0.1), with higher LC contrast associated to better sleep quality (**Figure 2B**) and no significant interaction between LC contrast and age group (p > 0.9)

**Figure 2.**
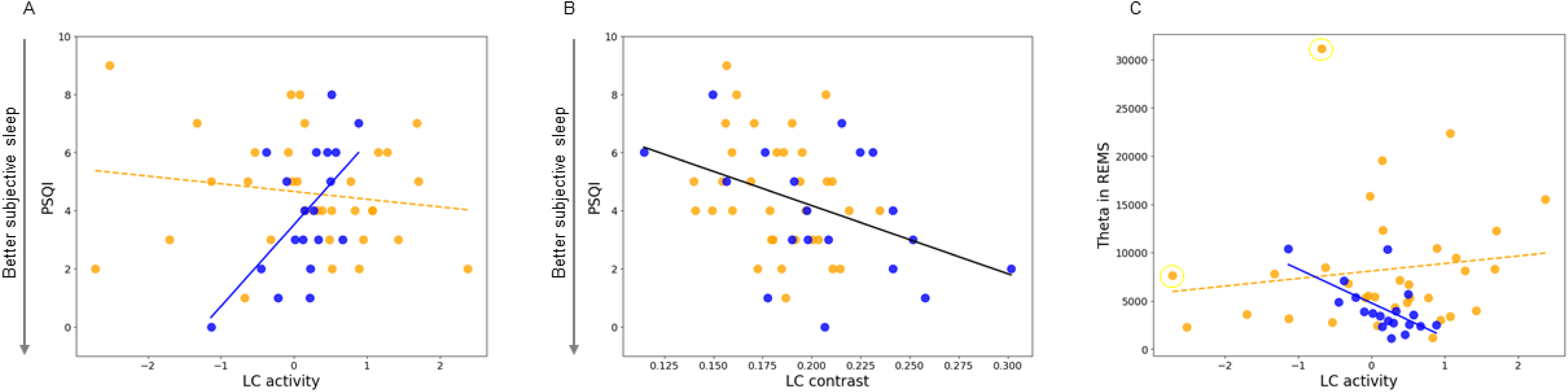
Associations between the LC and sleep metrics. (A) Association between habitual subjective sleep quality, as indexed by PSQI, and the activity of the LC. The GLMM yielded a significant age group by LC activity interaction and post-hoc analyses led to a significant association for the older but not the young group (cf. Table 2). (B) Association between habitual subjective sleep quality, as indexed by PSQI, and the LC contrast. The GLMM yielded a significant main effect of LC activity (cf. Table S1). (C) Association between the REMS theta power (cumulated overnight 4-8 Hz EEG power) and the LC activity with age-group interaction. The GLMM yielded a significant age group by LC activity interaction and post-hoc tests led to a significant association for the older but not the young group (cf. Table 2). The 2 highlighted dots correspond to two putative outliers [≥ 3 SD & < 4.5 SD for LC activity and REMS theta respectively] and we note that the p-value of the LC activity by age group interaction goes down to p = 0.012 when removed from the analyses. Orange dots represent younger individuals (18-30 y, N = 33) while the blue dots represent older individuals (50-70 y, N = 19). Simple regression lines are used for a visual display and do not substitute the GLMM outputs. The black line represents the regression irrespective of age groups (young + old, N=52). Solid and dashed regression lines are used for significant and non-significant outputs of the GLMM, respectively. The LC activity was computed as a mean of the activity estimates (betas) associated with the appearance of the target sounds in the bilateral LC mask of each subject, within the subject space. Displays are similar when using the left and right LC separately. Subjective sleep quality was estimated using the Pittsburgh Sleep Quality Index (PSQI) (Buysse, et al. 1989) where a higher score is indicative of some sleep difficulties.

**Table 2:**
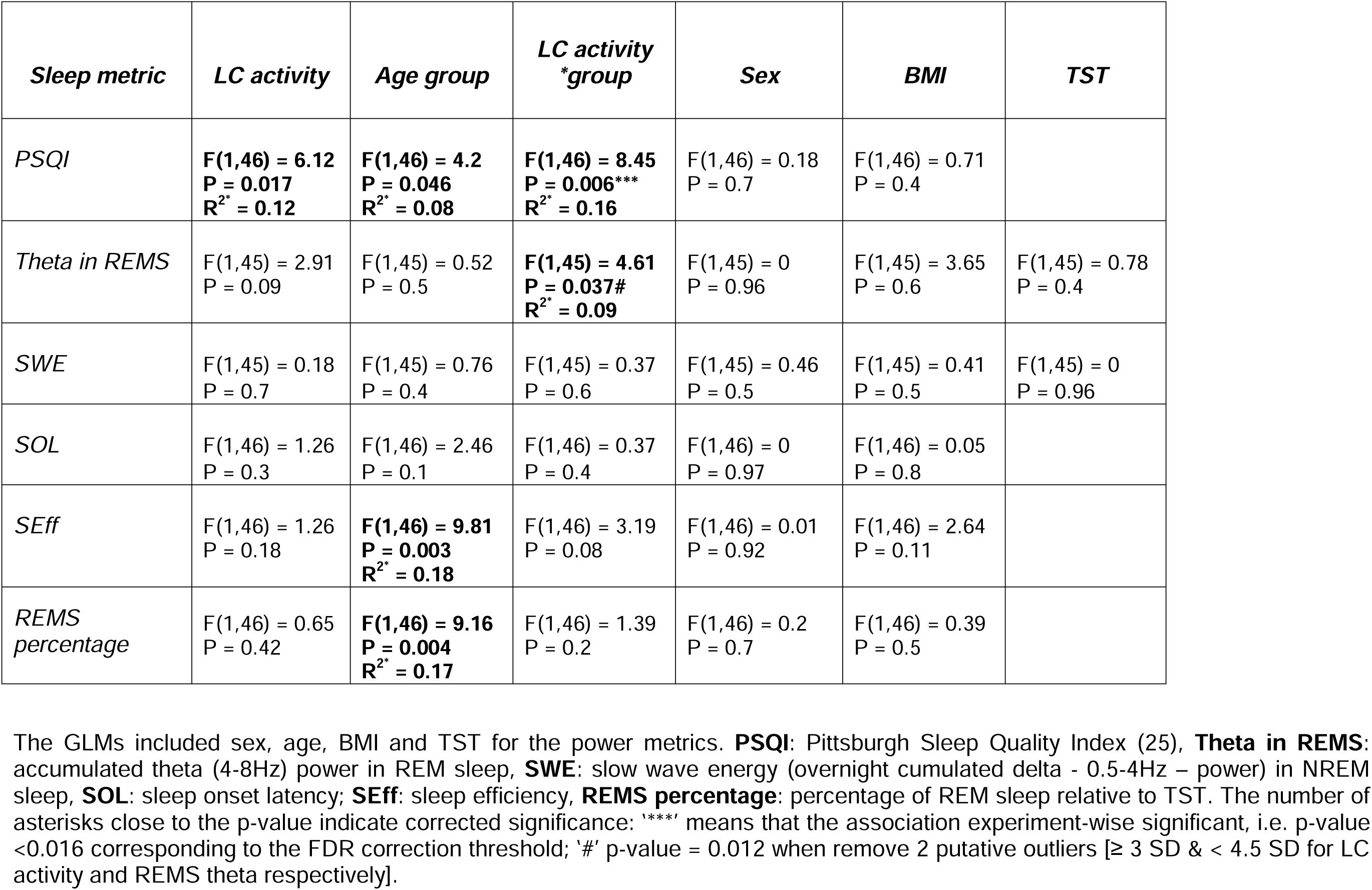
Statistical outcomes of GLMMs with the six baseline night sleep metrics of interest vs. the LC activity

We then considered the objective measures of sleep extracted from the EEG. A GLMM with the REMS theta power as the dependent variable found no main effect of the LC activity nor of age while it yielded a significant LC-activity-by-age-group interaction (**p = 0.037**), controlling for sex, BMI and total sleep time (TST) (**Table 2**). Post-hoc tests revealed that higher LC activity was associated with lower REMS theta power in the older (t = −2.02, **p= 0.049**), but not in the younger group (t= 0.81, p= 0.42) (**Figure 2C**). In addition, removing two putative outliers [≥ 3 SD & < 4.5 SD for LC activity and REMS theta respectively], the LC-activity-by-age-group interaction becomes even more robust (**p = 0.012;** **Figure 2C**). We then computed the same GLMM for mean activity of the left and right LC separately. The REMS theta power was significantly related to the activity of the left LC as a main effect (F = 4.49, **p= 0.04**, R^2*^ = 0.09) and there was an interaction with age (F = 5.33, **p = 0.026,** R^2*^ = 0.1), while no similar association was detected when using the activity of the right LC (F < 1.85, p > 0.15). Similar to the bilateral activity of the LC, post-hoc tests indicated that a higher activity of the left LC was related to lower REMS theta power in the older (t = −2.33, **p = 0.024**), but not in the younger group (t = 0.38, p = 0.7), while no similar association was found when focusing on the right LC (−1 < t < 1, p ≥ 0.3). As for subjective sleep quality, we added LC contrast to the GLMM which yielded the same LC-activity-by-age-group interaction, while no main effect of LC-contrast was detected (F = 0.05; p = 0.8) (**Table S1**). Lastly, if we control for REMS duration rather than for TST in the GLMM, statistical outputs lead to similar statistical tendencies (main effect of bilateral LC activity: p = 0.067; LC activity by group interaction: p = 0.055).

Importantly, none of the other sleep EEG metrics of interest were significantly associated with the activity of the LC (bilaterally or left and right separately) (**Table 2**), suggesting that the association was specific to the subjective sleep quality and REMS theta power. Given the close association between perceived sleep quality and REMS (28), we tested whether subjective sleep quality was correlated with the theta power in REMS in the older group, and we found a significant correlation with a large effect size (r = −0.54, **p = 0.016**) (**Figure 3A**), while the correlation was not significant in younger group and across the entire group (r = −0.26, p = 0.14; r = −0.22, p = 0.12). We further computed a mediation analysis that was purely exploratory given the size of the older subsample, to test whether the theta power in REMS mediated the association between the activity of the LC and subjective sleep quality in older individuals. The analyses yielded no statistical support for mediation (**Figure 3B**). While the direct link between LC activity and subjective sleep quality was significant (65.4 ± 32.8% of total effect; p = 0.046), the alternative indirect link was not significant (29.3 ± 41.4% of total effect; p = 0.48).

**Figure 3.**
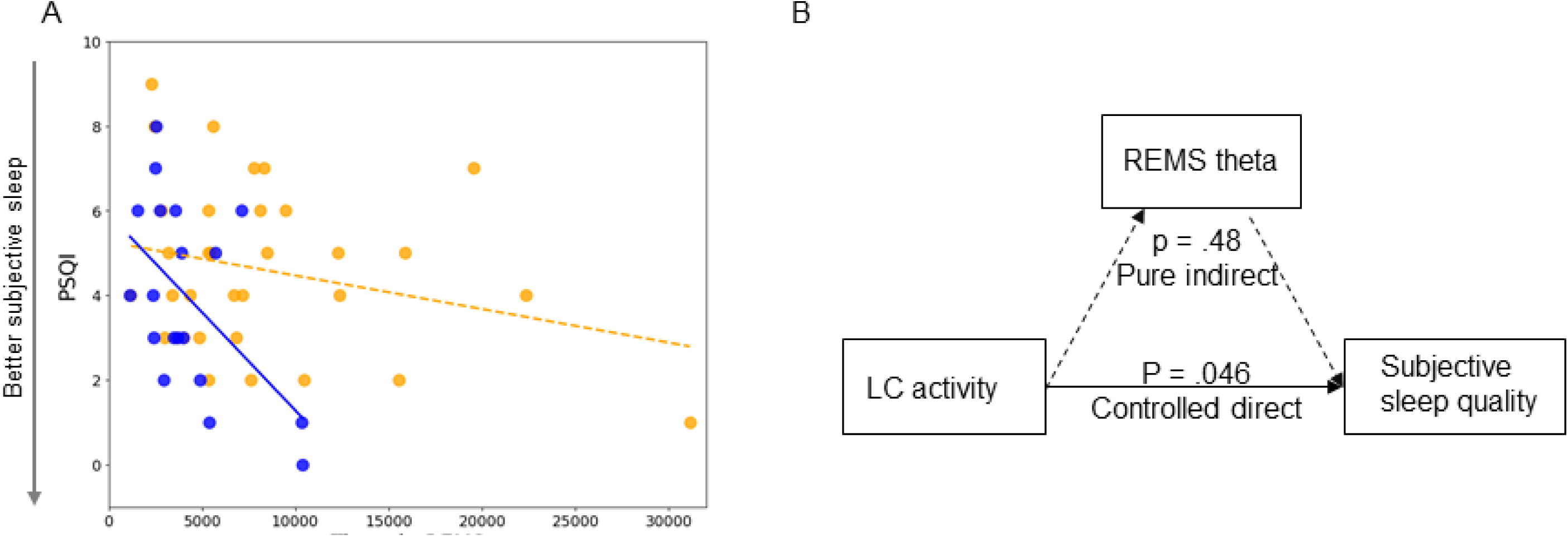
Associations between subjective sleep quality and REMS theta power. (A) Pearson’s correlation between habitual subjective sleep quality, as indexed by PSQI, and REMS theta power in the older group (N=19) (r = −0.54, P = 0.016). The orange dots represent individuals of the younger group (18-30 y, N = 33), and the blue dots represent individuals of the older group (50-70 y, N = 19). Solid and dashed regression lines are used for significant and non-significant Pearson’s correlations, respectively. (B) Mediation analyses in older individuals did not provide support for a mediation of the effect between the activity of the LC and subjective sleep quality by REMS theta power. The LC activity was computed as a mean of the activity estimates (betas) associated with the appearance of the target sounds in the bilateral LC mask of each subject, within the subject space. The LC contrast was computed as the mean contrast of the bilateral LC. Subjective sleep quality was estimated using the Pittsburgh Sleep Quality Index (PSQI) (Buysse, et al. 1989)

To gain further insight into the association between LC activity and REMS, we explored the association with additional REMS metrics, including the number of arousals during REMS and REMS episode duration. GLMMs with either of these metrics as the dependent variable did not lead to a significant main effect of LC activity (F ≤ 2.05; p ≥ 0.16) nor LC-activity-by-age-group interaction (F ≤ 3.03; p ≥ 0.08).

## Discussion

The LC is arguably one of the most important sleep-wake centers in the brain, and a growing number of animal and human studies have provided evidence supporting its role in regulating sleep and wakefulness, but that the precise mechanisms remain unknown. We provide evidence that, in contrast to young adults (18-30y), higher LC activity during wakefulness is associated with worse subjective sleep quality in late middle-aged individuals aged 50 to 70 y, cognitively unimpaired and devoid of sleep disorders. In addition, we show that higher LC activity during wakefulness is related to lower intensity of REMS in the older but not in the younger subsample. We further find that higher integrity of the LC, as indexed by the LC contrast, is associated with the better habitual subjective sleep quality across the entire sample.

*In vivo* recording of the activity of the LC during sleep is difficult. On top of its small size and deep position, the tonic activity of the LC is reduced and absent, respectively, during SWS and REMS, while generating sleep, and particularly REMS, in an MRI apparatus is not easy. However, fMRI is sensitive to individual variability in LC activity, whether during sleep or wakefulness, and one could argue that if the LC does not respond or activate as it should during a task, it might also indicate disrupted functional integrity and could be linked to impaired LC-modulated processes, such as sleep. In this first attempt to link LC function to the perceived quality of sleep and its electrophysiology *in vivo* in humans, we posit that the levels of activity of the LC during wakefulness and during sleep are directly related to one another. We decided to use a task known to induce a robust response of the LC during wakefulness (24) and link it with sleep features of interest.

With this in mind, the associations we find between LC activity, subjective sleep quality, and REMS intensity could arise from a negative impact of a higher activity of the LC during sleep (i.e. the higher activity we detected during wakefulness would “bleed” into sleep and correspond to a higher activity of the LC during sleep). In contrast to SWS, REMS quality, as indexed through the number of awakenings during REMS and the duration of REMS, constitutes a predictor of perceived sleep quality (28). One could therefore hypothesize that it is through the disturbance of REMS that the LC activity is associated with the perception of sleep quality. Although the significant correlation we find between the REMS theta power and the subjective sleep quality brings some support to this assumption, the mediation statistical analysis which we computed does not corroborate it. Future research should re-assess this mediation in a larger sample of old individuals.

Sleep quality begins to decline at age 40, and sleep complaints rise as well as adults get older (29,30). Many of the sleep alterations observed in aging and pathologies may arise from subcortical nuclei including the LC (13). We find that the association between the LC activity and sleep changes with age: higher LC activity is associated with poorer sleep quality and less intense REMS in middle-aged individuals and not in younger ones. Our findings support the idea that LC activity during sleep could shape part of the large inter-individual variability found in sleep disruptions, particularly starting at an age when sleep becomes more fragile, contributing therefore to age-related sleep complaints.

The LC modulates cortical activity through a tonic or phasic neuronal firing. A trade-off between these two modes allows for maximising the reward and the utility of incoming stimuli (31). Phasic bursts of LC-NE neurons are elicited when confronted with novel or salient stimuli such as in the oddball task we administered (21,32,33). Yet, our findings likely depend on the combination of tonic and phasic activity of the LC. Early LC damage has been suggested to result in a state of persistent high tonic LC activity that may disrupt task-related phasic activity (34). In addition, the temporal resolution of our fMRI data acquisition is relatively low compared to the burst of action potentials of LC (i.e., one volume was acquired in 2.34s). It is therefore hard to disentangle tonic and phasic contribution to our findings.

Interestingly, the LC, which is functionally connected to the salience network during wakefulness (35), presents an abnormal functional connectivity pattern in patients with insomnia disorder (36), which is the second most prevalent psychiatric disorder (37). This abnormal connectivity could contribute to the general state of hyperarousal characterizing insomnia disorder during both wakefulness and sleep to impede restful REMS (37). This assumption may underlie the association we find between LC activity and theta power during REMS. REMS theta activity is lower in patients with post- traumatic stress disorder, a condition often associated with insomnia, and higher REMS theta activity predicts a lower chance of re-experiencing symptoms following a stressful event (38). Theta oscillations during REMS are considered to be essential for the hippocampus-dependent memory consolidation during sleep (39), and they serve as the homeostatic control of REMS (40). Theta oscillations of REMS take place during a unique behavioral state when the LC is quasi-silenced, providing the conditions for the neuronal potentiation and de-potentiation required for a rewiring of the memory schemas depending on the hippocampus (37,41). Consequently, the negative association between LC activity and REMS theta power we find could reflect a relatively more restless REMS when the LC is insufficiently silenced with potential disruption in synaptic plasticity and memory consolidation (42). REMS would therefore be maladaptive to the dissolving of distress leading to a higher level of general anxiety. Here, we probed LC activity using an oddball task known to recruit the salience network (27) and we used a sample of individuals devoid of sleep and anxiety disorders. Hence, our findings could consist of the healthy spectrum of the association between LC activity and REMS that would lead to insomnia disorder if exacerbated or prolonged over extended periods of time. Future investigations are warranted in a clinical population with, for instance, anxiety and/or insomnia disorder, including tests of memory performance as well as other behavioral measures.

We fortuitously find that a higher LC MRI contrast is associated with better subjective sleep quality. This finding echoes a recent report that lower LC contrast in its middle-caudal portion is linked to a higher number of self-reported nocturnal awakenings in healthy older individuals (19). Although the LC contrast is considered to reflect its structural integrity, its neurobiological bases are still under investigation (43). The LC contrast increases over the adult lifespan up to around 55-60y to decrease after (44), preventing our understanding on whether a higher LC contrast over time reflects a better or worse situation. In our sample of very healthy individuals, the LC is possibly better preserved (e.g. it may present less tau aggregates (45)), such that higher contrast is associated with better sleep quality.

According to autopsy data, by the age of 40y, about 100% of the population exhibits some degree of tau protein aggregates in the LC (15). The presence of these tau aggregates is likely to affect the LC structure and functioning (46), and is suspected to contribute to cognitive decline in older individuals (47). Since we did not assess tau aggregates levels, we cannot address whether tau aggregates contribute to our findings. Similarly, our findings may suggest more prominent associations between the left LC and sleep, however, we do also find associations with the right LC. Since there is no clear consensus on the potential lateralization of the LC (12,48), we have reported here a potential laterality while we do not interpret our findings in terms of lateralization.

Our study has limitations. Most importantly, the timeline of the experiment was different between the younger and older participants. While the baseline night of sleep immediately preceded the fMRI acquisition in the young, a time gap of up to 1 year separates these 2 parts in the older group (see methods). Although sleep undergoes profound changes over the lifetime (49), it is remarkably stable within an individual over a shorter period (e.g. a few months/years) (50). While we acknowledge that the difference in procedure can induce bias, we consider it unlikely to explain the significant association we find in the older, but not in the younger individuals. Our sample also included a larger proportion of woman (73 to 85%) such that differences between sexes, though accounted for in the statistical modeling, could not be properly investigated. Our sample size, while representing a large work effort, remains modest, particularly in the older subsample, and replication is warranted. Future studies may also want to use individually tailored hemodynamic response functions (HRF) to assess LC response. While the canonical HRF we used to model activity over the entire brain seems suitable to model average LC response over a group of participants, individual LC responses can vary substantially across individuals (51). Finally, applying an False Discovery Rate (FDR) correction to the p-values of the primary GLMMs, only the association between LC activity and sleep quality remains significant (interaction with the age group) while the association with REMS theta power does not reach the corrected p-value threshold (i.e. p < 0.016, see methods), though it may meet the corrected threshold if subject with relatively extreme values are removed. Given that our study is pioneering in seeking relations between sleep features and LC activity during wakefulness *in vivo* in humans, the value of our results remains highly remarkable.

We provide *in vivo* evidence that higher LC function is associated with both worse subjective and objective measures of sleep quality in healthy individuals, particularly in the late middle-aged group. Sleep complaints and alterations in the regulation of sleep show higher prevalence with ageing and constitute strong risk factors for the development of insomnia disorder (37). In addition, age-related alterations in the regulation of sleep, including REMS, bear some predicting values for the future development of AD (52,53). Our results contribute to the understanding of the physiology of sleep and may therefore also have implications for the treatment of clinical populations that would target sleep or the LC (37).

## Materials and Methods

### Participants

A sample of 52 healthy participants of both sexes, composed of 33 healthy young (age: 22.15 ± 3.27 y, 28 women) and 19 late middle-aged (age: 61.05 ± 5.3 y, 14 women) individuals were recruited from the local community to participate to this study. A summary of the demographic data can be found in Table 1. This study was approved by the faculty-hospital ethics committee of the University of Liège. All participants provided their written informed consent and received financial compensation.

The exclusion criteria were as follows: history of major neurologic or psychiatric diseases or stroke, a recent history of depression and anxiety, sleep disorders, medication affecting the central nervous system, smoking, excessive alcohol (> 14 units/week) or caffeine (> 5 cups/day) consumption, night shift work in the past 6 months, travel across different time zones during the last 2 months, Body Mass Index (BMI) ≤ 18 and ≥ 29 (for the older participants) and ≥ 25 (for the younger participants), clinical symptoms of cognitive impairment for older subjects (dementia rating scale score < 130; Mini-Mental State Examination score < 27) and MRI contraindications. Due to a miscalculation at screening one older participant had a BMI of 30.9 and one of the younger participants had a BMI of 28.4. Since their data did not deviate substantially from the rest of the sample and BMI was used as a covariate in our statistical models, these participants were included in the analyses.

### Protocol

All participants completed an in-lab habituation night under polysomnography to minimize the effect of the novel environment for the subsequent baseline night and to exclude volunteers with sleep disorders (see below). All participants completed a whole-brain structural MRI acquisition and an acquisition centered on the LC using a specific sequence. They further completed the PSQI questionnaire to assess their habitual subjective sleep quality (25). Higher scores are indicative of some sleep difficulties. Participants were requested to sleep regularly for 7 days before the baseline night (±30 min) based on their preferred bed and wake-up times. The compliance was verified by the sleep-wake diary and actigraphy (Actiwatch©, Cambridge Neurotechnology, UK and AX3©, Axivity Ltd, UK). Participants were instructed to abstain from caffeinated beverages and from alcohol as well as to not do excessive physical activity at least three days before the baseline night. The evening before the baseline night participants arrived at the laboratory 4 hours before their habitual bedtime, completed questionnaires including the Beck Anxiety Inventory (BAI) and Beck Depression Inventory (BDI) (54,55), and were then kept in dim light (<10 lux) for 3 hours preceding bedtime. Their habitual sleep was then recorded in complete darkness under EEG. Baseline night data was acquired using N7000 amplifiers (EMBLA, Natus Medical Incorporated, Planegg, Germany) and were used for sleep features extraction. All participants completed a fMRI session that included three perceptual tasks, approximately 3 hours after habitual wake-up time. This paper is centered on the analyses of the oddball auditory task brain responses.

Younger participants completed the fMRI session immediately following the in-lab baseline night. They were maintained in dim light (< 10 lux) between wake-up time and the fMRI session. Older participants were part of a different study (56,57) and completed the sMRI and fMRI procedures in addition to their initial engagement. Consequently, the baseline nights of sleep and fMRI sessions were completed about 1 year apart (mean ± SD: 15.5 ± 5.3 months). The procedures for the baseline night recordings, including the sleep-wake schedule and light exposures, were identical to those of the young. Prior to the fMRI session participants slept regularly for 1 week (verified with a sleep diary; our experience is that actigraphy reports and sleep diaries do not deviate substantially in older individuals); they were maintained in dim light (< 10 lux) for 45 min before the fMRI scanning.

### Sleep EEG metrics

The habituation night included 5 EEG derivations (Fz, Cz, Pz, P3, Oz) while 11 derivations were used for the baseline night (F3, Fz, F4; C3, Cz, C4; P3, Pz, P4; O1, O2), all placed according to the 10–20 system and referenced to the left mastoid (A1) while an electrode was also placed over the right mastoid (A2). Both nights included 2 bipolar electrooculogram (EOG), and 2 bipolar submental electromyogram (EMG) electrodes as well as 2 bipolar electrocardiogram (ECG) derivations. EEG data was digitized at 200 Hz sampling rate. EEG data was then re-referenced off-line to the average of both mastoids using Matlab (Mathworks Inc., Sherbom, MA). Participants with excessive sleep apneas (apnea-hypopnea index ≥ 15) and limb movements (and limb movements (≥ 15/hour) were excluded from the study following the habituation night. No participants suffered from REMS behavioral disorder nor from other parasomnia.

The sleep data were scored in 30-s epochs using a validated automatic sleep scoring algorithm (ASEEGA, PHYSIP, Paris, France) (58). Arousals and artefacts were detected automatically as previously described (59) and excluded from the subsequent power spectral density analyses (using Welch’s overlapped segment averaging estimator, as implanted in the pwelch Matlab function; 4s epochs without artefact or arousal; 2s overlap). Only frontal electrodes were considered in the analyses because the frontal region is most sensitive to sleep pressure manipulations (60) and to facilitate interpretation of future large-scale studies using headband EEG, often restricted to frontal electrodes. Averaged power was computed per 30 min bins, adjusting for the proportion of rejected data (containing artefact/arousal), and subsequently aggregated in a sum separately for REM and NREM sleep as described in (61). Thus, we computed slow wave energy (SWE) - cumulated power in the delta frequency band during SWS, an accepted measure of sleep need (26), and, similarly, we computed the cumulated theta (4-8Hz) power in REM sleep. The cumulated power score would increase with time spent in REMS and SWS, so we included TST as a common covariate in all analyses, as well as REMS duration in secondary analyses.

### Auditory oddball task

The task consisted of rare deviant target tones (1000 Hz sinusoidal waves, 100 ms), composing 20% of the tones that were pseudo-randomly interleaved within a stream of standard stimuli (500 Hz sinusoidal waves, 100 ms). The task included 270 auditory stimuli in total (54 target tones). Auditory stimuli were delivered with MRI-compatible headphones (Sensimetrics, Malden, MA). The inter-stimulus interval was set to 2000 ms. Participants were instructed to press with the right index on an MRI-compatible keyboard (Current Designs, Philadelphia, PA) as quickly as possible at the appearance of target sounds. The experimental paradigm was designed using OpenSesame 3.3.8 (Mathôt et al., 2012). The MRI session started with a short session to set the volume of the audio system to ensure an optimal perception of the stimuli.

### MRI data acquisitions

MRI data were acquired using a MAGNETOM Terra 7T MRI system (Siemens Healthineers, Erlangen, Germany), with a single-channel transmit coil and a 32-receiving channel head coil (1TX/32RX Head Coil, Nova Medical, Wilmington MA). To reduce dielectric artifacts and homogenize the magnetic field of Radio Frequency (RF) pulses, dielectric pads (Multiwave Imaging, Marseille, France) were placed between the head of the participants and the coil.

BOLD fMRI data were acquired during the task, using a multi-band Gradient-Recalled Echo - Echo-Planar Imaging (GRE-EPI) sequence: TR = 2340 ms, TE = 24 ms, flip angle = 90°, matrix size = 160 × 160, 86 axial 1.4mm-thick slices, MB acceleration factor = 2, GRAPPA acceleration factor = 3, voxel size = (1.4×1.4×1.4) mm^3^. The cardiac pulse and the respiratory movements were recorded concomitantly using, respectively, a pulse oximeter and a breathing belt (Siemens Healthineers, Erlangen, Germany). The fMRI acquisition was followed by a 2D GRE field mapping sequence to assess B0 magnetic field inhomogeneities with the following parameters: TR = 5.2 ms, TEs = 2.26 ms and 3.28 ms, FA = 15°, bandwidth = 737 Hz/pixel, matrix size = 96 × 128, 96 axial slices, voxel size = (2×2×2) mm ^3^, acquisition time = 1:38 min.

A Magnetization-Prepared with 2 RApid Gradient Echoes (MP2RAGE) sequence was used to acquire T1 anatomical images: TR = 4300 ms, TE = 1.98 ms, FA = 5°/6°, TI = 940 ms/2830 ms, bandwidth = 240 Hz/pixel, matrix size = 256 × 256, 224 axial 0.75mm-thick slices, GRAPPA acceleration factor = 3, voxel size = (.75x.75x.75) mm^3^, acquisition time = 9:03 min (Marques & Gruetter, 2013). The LC specific sequence consisted of a 3D high-resolution Magnetization Transfer-weighted Turbo-FLash (MT-TFL) sequence with the following parameters: TR = 400 ms, TE = 2.55 ms, FA = 8°, bandwidth = 300 Hz/pixel, matrix size = 480 × 480, number of averages = 2, turbo factor = 54, MTC pulses = 20, MTC FA = 260°, MTC RF duration = 10000 µs, MTC Inter RF delay = 4000 µs, MTC offset = 2000 Hz, voxel size = (.4x.4x.5)mm^3^, acquisition time = 8:13 min. Sixty axial slices were acquired and centered for the acquisitions perpendicularly to the rhomboid fossa (i.e., the floor of the fourth ventricle located on the dorsal surface of the pons).

### MRI data pre-processing

EPI images underwent motion correction, distortion correction using Statistical Parametric Mapping (SPM12, https://www.fil.ion.ucl.ac.uk/spm/) and brain extraction using “BET” from the FMRIB Software Library suite (FSL, https://fsl.fmrib.ox.ac.uk) and the final images were spatially smoothed with a Gaussian kernel characterized by a full-width at half maximum of 3 mm.

The background noise in MP2RAGE images was removed using an extension of SPM12 (extension: https://github.com/benoitberanger/mp2rage) (62). The denoised images were then automatically reoriented using the ‘spm_auto_reorient’ function (extension available from https://github.com/CyclotronResearchCentre/spm_auto_reorient) and corrected for intensity non-uniformity using the bias correction method implemented in the SPM “unified segmentation” tool (63). Brain extraction was then conducted on the denoised-reoriented-biased-corrected image using both the Advanced Normalization Tools (ANTs, http://stnava.github.io/ANTs/) with the ‘antsBrainExtraction’ function and the RObust Brain EXtraction tool (ROBEX, https://www.nitrc.org/projects/robex) (64). The method yielding to the best extraction for each individual, as assessed by visual inspection, was used for subsequent steps. A whole-brain T1 group template was created using ANTs, based on preprocessed MP2RAGE images of all subjects except for one, the MP2RAGE image of whom was not suitable due to a bad positioning of the field of view during the acquisition. Finally, the preprocessed MP2RAGE image of each subject was normalized to the Montreal Neurological Institute (MNI) space (with a 1×1×1 mm^3^ image resolution). The purpose of using a template that is specific to our dataset was to improve the registration into the MNI space using a study specific intermediate space. The transformation parameters obtained from normalization were later used for registering first-level statistical maps into the MNI space to conduct group-level fMRI analyses.

To extract the LC contrast, T1 structural images in subject space (after removing the background noise) were up-sampled by a factor 2 [(0.375×0.375×0.375) mm^3^] to avoid losing in-plane resolution when registering the LC slab to the T1 image. The up-sampling was done using the ‘nii_scale_dims’ function from an extension of SPM12 (extension: https://github.com/rordenlab/spmScripts). The complete LC contrast extraction was done in the original subject space. The MT-TFL image of each subject was registered with the whole brain up-sampled T1 image by means of a two-step process: (i) an approximate manual registration to extract the parameters for an initial transformation using ITK-SNAP (65), and (ii) an automatic affine registration based on the initial transformation parameters, using ANTs. MT-TFL data of one young subject was not usable, due to the excessive motion of the participant, leading to a registration failure.

The LC appearing hyperintense on registered MT-TFL images was manually delineated by two expert raters and the intersection of the LC masks of the two raters was computed as the final LC mask for each individual. The LC mask was skeletonized by only keeping the voxel with the highest intensity in each axial slice. Based on the skeletonized LC mask, the LC contrast was computed after normalization of each LC slice intensity to a slice-corresponding 2D reference region (a 15 x 15 voxels region, corresponding to a (5.5 x 5.5)mm^2^ square region) situated anteriorly (and centrally) in the pons, in the pontine tegmentum. For example, the left LC contrast was defined as:

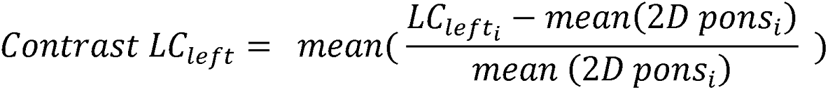

Where:

- i is the slice index along the (left) LC,
- *LC_Left_*, is the intensity of the voxel with the highest intensity in the axial slice with index i,
- mean (2D pons_i_) represents the mean intensity in the 2D reference region corresponding to the axial slice with index i.

The LC contrast was computed as the mean LC contrast between the left and right LC. The distribution of the LC contrasts across individuals was investigated by computing the Probability Density Function (PDF), using a kernel density nonparametric method (ks density MATLAB R2021a built-in function). Individual skeletonized LC masks were used for extracting the LC activity during the oddball task in the subject space. To investigate the activation of the LC at the group level, an LC group-wise template was created. The LC mask of each volunteer was normalized to the structural group template, and then to the MNI (MNI152 - 1×1×1mm^3^). This was done using the ‘antsApplyTransforms’ ANTs command, with the transformation parameters estimated (i) when registering the subject-specific MP2RAGE image to the structural template, and (ii) the transformation parameters estimated when registering the structural template to the MNI. The final LC group-wise template was created as the sum of all masks.

## Statistical Analyses

Statistical analyses were conducted using SPM12. A high-pass filter with a 128 s cutoff was applied to remove slow signal drifts. The timing vector with the appearance of the target tones was convolved with the canonical HRF to model the event-related response and was used as the main condition in a General Linear Model (GLM). The PhysIO Toolbox (https://www.tnu.ethz.ch/en/software/tapas/documentations/physio-toolbox) was used to compute physiology-related voxel-wise signal fluctuations based on respiratory and cardiac pulsation data (66), that was available in 48 volunteers (physiological data was not available for 4 volunteers). The Fourier expansion of cardiac and respiratory phase, 14 parameters computed with the toolbox, as well as the 6 realignment parameters were used as multiple regressors of no interest in the GLM. The first-level statistical analysis was conducted in the subject space.

The mean functional image was registered to the MP2RAGE image to extract the corresponding transformation matrix used to register the first-level statistical map of each subject to the structural image. Therefore, for all subjects, statistical maps corresponding to the appearance of target sounds were registered to the native space, normalized to the group template space and then to the MNI space. A second-level analysis was then conducted in the MNI space, where age, sex and BMI were used as covariates. The group-wise mask of the LC was used to assess specific activation of the LC. Due to the small size of the nucleus, LC activation was not expected to survive stringent whole-brain FWE correction. Therefore, a small volume correction using the LC template was conducted using SPM12 to detect voxel-level p<.05 FWE-corrected results within the LC mask.

REX Toolbox (https://web.mit.edu/swg/software.htm) was then used to extract the activity estimates (betas) associated with the appearance of the target sounds in the LC mask of each subject, within the subject space (67). This procedure ensured that any potential displacement and bias introduced by the normalization step into the common MNI space did not affect individual activity estimates. Statistical analyses using these activity estimates were performed in SAS 9.4 (SAS Institute, NC, USA). Analyses consisted of Generalized Linear Mixed Model (GLMM) with sleep features of interest as the dependent variable, the LC activity as an independent variable and age group (younger, older), sex, BMI included as covariates, and subject as a random factor. Partial R^2^ (R^2*^) values were computed to estimate the effect sizes of significant effects in each model (68). GLMM were computed according to the distribution of the dependent variable. In the primary analyses, we tested 6 independent GLMMs, and to account for multiple comparisons, we applied the Benjamini–Hochberg procedure for false discovery rate correction of the p-values using and online tool https://tools.carbocation.com/FDR. The tool yielded a corrected p-value of 0.016.

The mediation analysis was computed using CAUSALMED procedure in SAS including bootstrap confidence interval computation. Subjective sleep quality was the dependent variable with a direct pathway to LC activity and an indirect pathway mediated by theta power in REMS, which was squared root to satisfy the parametric assumption of the procedure. An interaction effect between theta power in REMS and LC activity was included, while age, sex and BMI were used as covariates. The percentage of controlled direct and pure indirect effects are reported together with their associated p-values.

Optimal sensitivity and power analyses in GLMMs remain under investigation (e.g. (69)). We nevertheless computed a prior sensitivity analysis to get an indication of the minimum detectable effect size in our main analyses given our sample size. According to G*Power 3 (version 3.1.9.4) (70), taking into account a power of 0.8, an error rate α of 0.05, and a sample size of 52 (33 + 19), we could detect medium effect sizes r > 0.33 (one-sided; absolute values; CI: 0.06–0.55; R²>0.11, R² CI: 0.003–0.3) within a linear multiple regression framework including one tested predictor (LC activity) and three/four covariates (group, sex, BMI, and TST where relevant).

## Data availability

The data and analysis scripts supporting the results included in this manuscript are publicly available via the following open repository: https://gitlab.uliege.be/CyclotronResearchCentre/Public/xxx *(to be done following peer reviewing and upon acceptance for publication / and editor request)*. We used Matlab, bash and/or Python scripts for EEG and MRI data processing, while we used SAS for statistical analyses. Researchers willing to access the raw data should send a request to the corresponding author (GV). Data sharing will require evaluation of the request by the local Research Ethics Board and the signature of a data transfer agreement (DTA).

## Acknowledgements

We thank N Beliy, E Lambot, C Hagelstein, S Laloux, A Claes, C Degueldre, B Herbillon, G Hammad, P Hawotte, B Lauricella and A Lesoinne for their help in the different steps of the study.

This work was supported by Fonds National de la Recherche Scientifique (FRS-FNRS, T.0242.19 & J. 0222.20). Action de Recherche Concertée – Fédération Wallonie-Bruxelles (ARC SLEEPDEM 17/27-09), Fondation Recherche Alzheimer (SAO-FRA 2019/0025), University of Liège (ULiège), European Regional Development Fund (Radiomed & Biomed-Hub). AB is supported by Synergia Medial SA and the Walloon Region (Industrial Doctorate Program, convention n°8193). EB is supported by the Maastricht University - Liège University Imaging Valley. RS and FB are supported by the European Union’s Horizon 2020 research and innovation program under the Marie Skłodowska-Curie grant agreement No 860613. EK, IP, IC, NM, CP, KK, NM, GV are supported by the FRS-FNRS. SS was supported by ULiège-Valeo Innovation Chair and Siemens Healthineers. HJ is supported by the National Institutes of Health [R01AG062559, R01AG068062, R21AG074220]; the Alzheimer’s Association [AARG-22-920434] and Alzheimer Nederland major award [WE.03-2019-02].

---

**Table S1:**
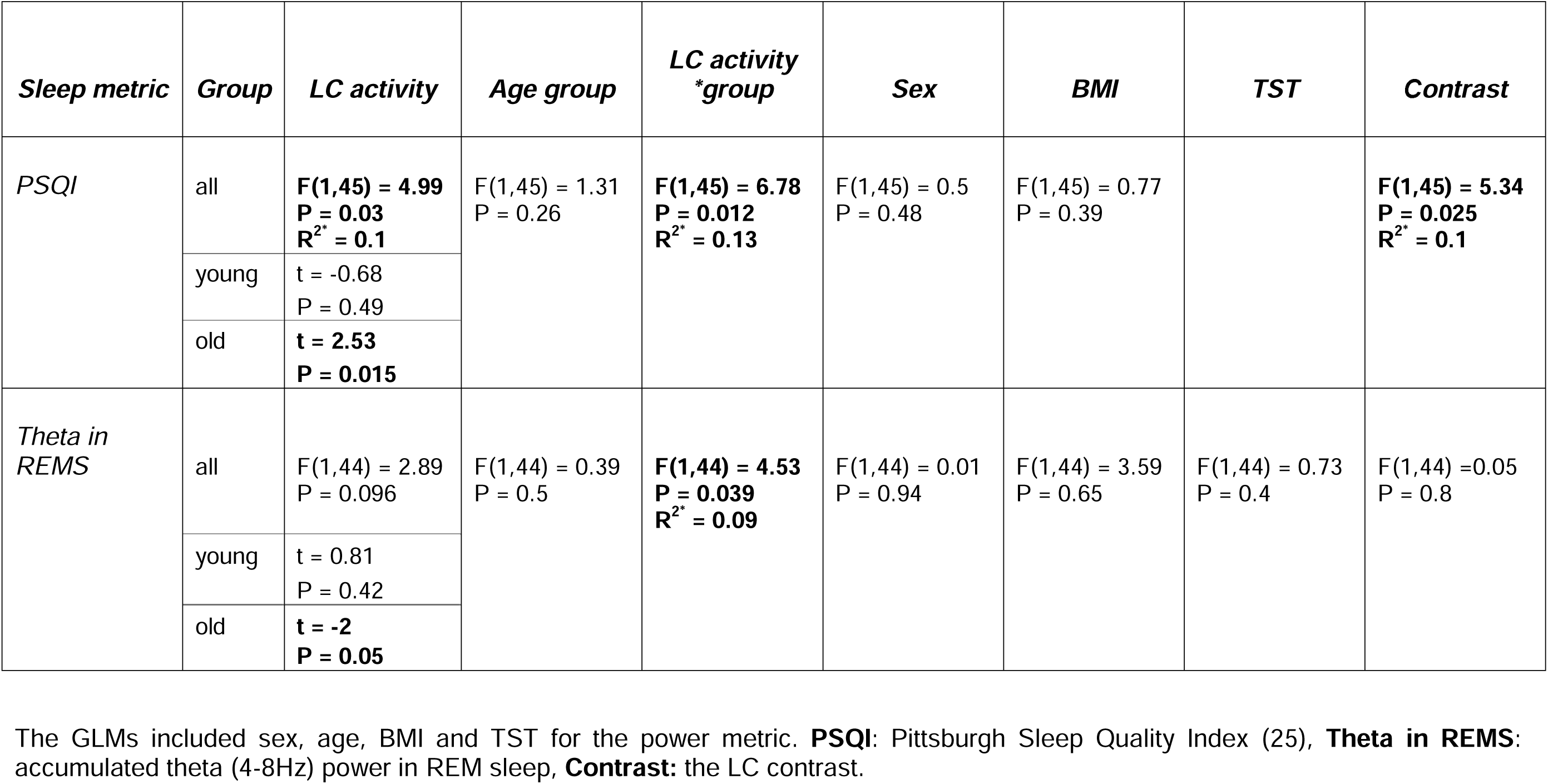
Statistical outcomes of GLMMs with the two sleep metrics of interest vs. the LC activity while controlling for the LC contrast

| *Sleep metric* | *Group* | *LC activity* | *Age group* | *LC activity*<br>*\*group* | *Sex* | *BMI* | *TST* | *Contrast* |
| --- | --- | --- | --- | --- | --- | --- | --- | --- |
| *PSQI* | all | **F(1,45) = 4.99**<br>**P = 0.03**<br>**R<sup>2</sup>\* = 0.1** | F(1,45) = 1.31<br>P = 0.26 | **F(1,45) = 6.78**<br>**P = 0.012**<br>**R<sup>2</sup>\* = 0.13** | F(1,45) = 0.5<br>P = 0.48 | F(1,45) = 0.77<br>P = 0.39 |  | **F(1,45) = 5.34**<br>**P = 0.025**<br>**R<sup>2</sup>\* = 0.1** |
|  | young | t = -0.68<br>P = 0.49 |  |  |  |  |  |  |
|  | old | **t = 2.53**<br>**P = 0.015** |  |  |  |  |  |  |
| *Theta in*<br>*REMS* | all | F(1,44) = 2.89<br>P = 0.096 | F(1,44) = 0.39<br>P = 0.5 | **F(1,44) = 4.53**<br>**P = 0.039**<br>**R<sup>2</sup>\* = 0.09** | F(1,44) = 0.01<br>P = 0.94 | F(1,44) = 3.59<br>P = 0.65 | F(1,44) = 0.73<br>P = 0.4 | F(1,44) = 0.05<br>P = 0.8 |
|  | young | t = 0.81<br>P = 0.42 |  |  |  |  |  |  |
|  | old | **t = -2**<br>**P = 0.05** |  |  |  |  |  |  |

The GLMs included sex, age, BMI and TST for the power metric. **PSQI**: Pittsburgh Sleep Quality Index (25), **Theta in REMS**: accumulated theta (4-8Hz) power in REM sleep, **Contrast**: the LC contrast.

